# The central THαβ immunity associated cytokine: IL-10 has a strong anti-tumor ability toward established cancer models in vivo and toward cancer cells in vitro

**DOI:** 10.1101/2020.10.19.344804

**Authors:** Wan-Chung Hu

## Abstract

Cancer immunotherapy is a promising new approach for cancer treatment. We propose to use a new host immunological pathway: THαβ immunity for cancer treatment. THαβ immunity is a natural host immunity which can be used IFN-α/β and its major effectors are IL-10 and IL-15. Thus, we here used these cytokines for treatment both in a mice breast cancer 4T1 model and in a neuroblastoma NXS2 model. 4T1 cells and NXS2 cells were inoculated subcutaneously to wild type BALB/c female mice and AJ mice, respectively. Both Cytokines and an antibody treatment with variable dosages were given. Our results show that IL-10 and IL-15 treatments have significant effects on not only tumor volume shrinkage and but also on mice survival. However, IFN-γ even worsens the tumor volume and mice survival. Antibodies plus IL-10 is no better than IL-10 alone for cancer immunotherapy. This can be due to GD2 antigen expression on immune cells. Moreover, an anti-GD2 antibody could harm immune cells themselves. We found that IL-10 has direct toxicity on tumor cells in vitro. Our results conclude that using THαβ immunity is a good strategy for cancer immunotherapy.

## Introduction

Current cancer treatments, including surgery, radiotherapy, and chemotherapy are not satisfactory. Immunotherapy is a prospective new treatment strategy for cancer. Current immunotherapy using monoclonal antibodies are effective. However, resistance to monoclonal antibody treatment has been reported during cancer therapy. Thus, we will need better ways to activate host immunological pathways against cancer. Cytokines are important players in initiating immune cells. Thus, we can use cytokines as immunostimulants during monoclonal antibodies treatment against cancer.

Host immunological pathways can be divided into four branches(1, 2). TH2 is an immunity response that against helminths. Its effectors are eosinophils, basophils, mast cells, IgE/IgG4 secreting B calls, and IL4/IL5 secreting CD4 T cells. TH17 is an immunity response against extracellular bacteria. Its effectors are neutrophils, IgA/IgG2 secreting B cells, and TGFβ/IL6/IL17 secreting CD4 T cells. Traditional TH1 is an immunity response used against intracellular bacteria. Its effectors are macrophages, IgG3 secreting B cells, cytotoxic T cells, and IFN-γ secreting CD4 T cells. In my previous study, I proposed a new host immunological pathway THαβ immunity. THαβ immunity is an immunity response against viruses. Its effectors are NK cells, IgG1 secreting B cells, cytotoxic T cells, and IL10/IFN-α/β secreting CD4 T cells(3). On this study, we used THαβ immunity as a cancer immunotherapy. In THαβ immunity, NK cell antibody dependent cellular cytotoxicity (ADCC) is activated to kill virus infected cells by means of NK cells’ binding to cell surface viral antigen. In this research, we investigated cytokine adjuvants’ role for activating NK cell ADCC mechanism.

Here, we used a 4T1 mouse breast cancer cell line and an NXS2 mouse neuroblastoma cell line as our tumor in vivo models(4). 4T1 cells were inoculated subcutaneously in Balb/c female mice to set up the animal model. Cytokines (IFN-α/β, IL-10, IL-15, and IFN-γ) and monoclonal anti-Globo H antibody (VK9) were injected intraperitoneally to mice for 5 days(5). We find that IL-15 and IL-10 have significant effects which cause tumor volume shrinkage and prolong mice survivals. In the second NXS2 cell AJ mice model, IL-10, anti-GD2 14G2A antibody, combined with IL10 plus an anti-GD2 antibody were used for cancer immunotherapy(6, 7). We found that only IL-10 alone is the most effective treatment strategy. We conclude that THαβ immunity is effective for cancer immunotherapy.

## Results

### IL-10 has had significant effects on tumor shrinkage and mouse survival in the 4T1 model

Our results show that IL10 (20ug IP x 5days) has had significant results with respect to both tumor volume shrinkage and mouse survival (Figure 1). After the IL-10 injection, tumor size is much smaller compared with that of the control mice. In addition, IL-10 treatment has prolonged the mice survival. Our findings suggested that IL-10 is potentially a very good candidate for cancer immunotherapy.

**Figure 1-1:**
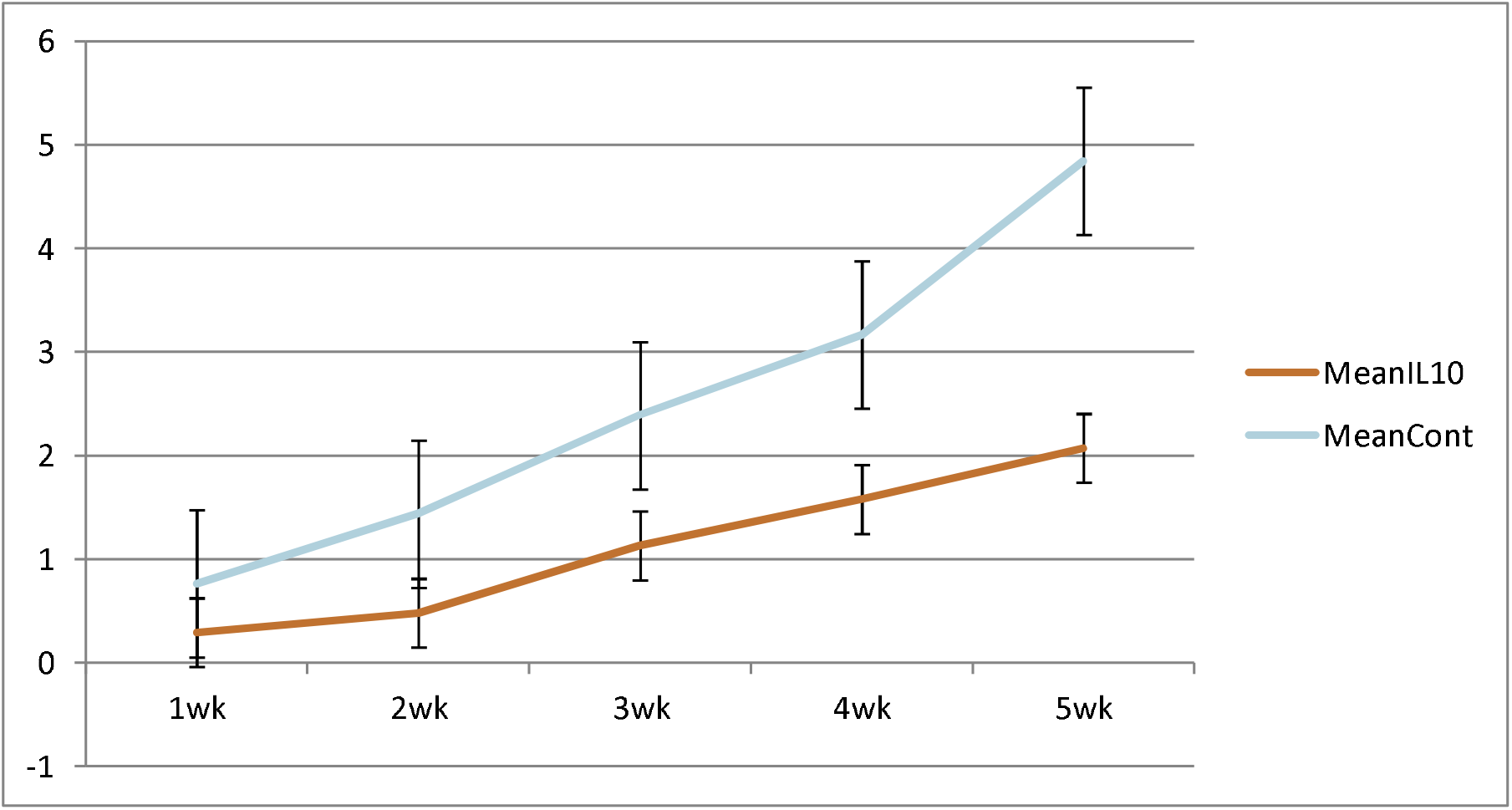
Tumor volume was measured every week after 4T1 cell inoculation with or without IL-10 treatment. Blue line means control group and red line means the IL-10 group. IL-10 therapy caused statistically significant tumor volume shrinkage(standard error is used in the graph)

**Figure 1-2:**
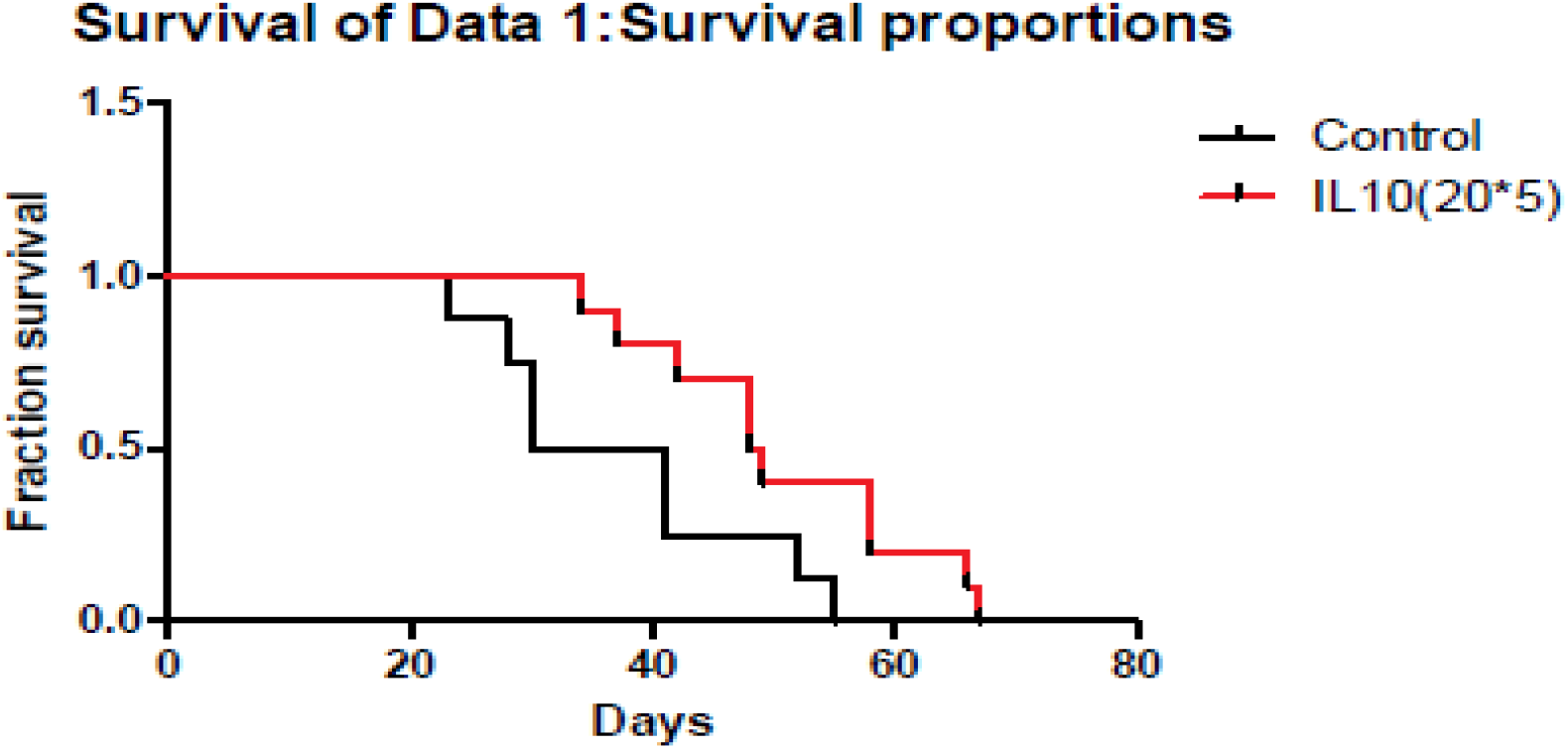
Mice survival was measured each week after 4T1 cell inoculation with or without IL-10 treatment. Red line means IL-10 treated group and black line means control group. IL-10 treatment significantly prolonged mice survival. (Likelihood P value=0.020, median survival for control group=35.5 days and for IL-10 treated group=48.5 days)

### IL-15 decreases tumor volume and prolongs mice survival in the 4T1 model

Cytokine IL-15 also can suppress tumor growth and prolong mice survivals (Figure 2). After IL-15 treatment (15ug bid IP x 5days), the tumor volume is significantly smaller compared to that of the control mice, especially in the late stage. IL-15 also prolongs mice survivals. After IL-15 injection, mice inoculated with 4T1 cells survived 2 weeks more than the control mice.

**Figure 2-1:**
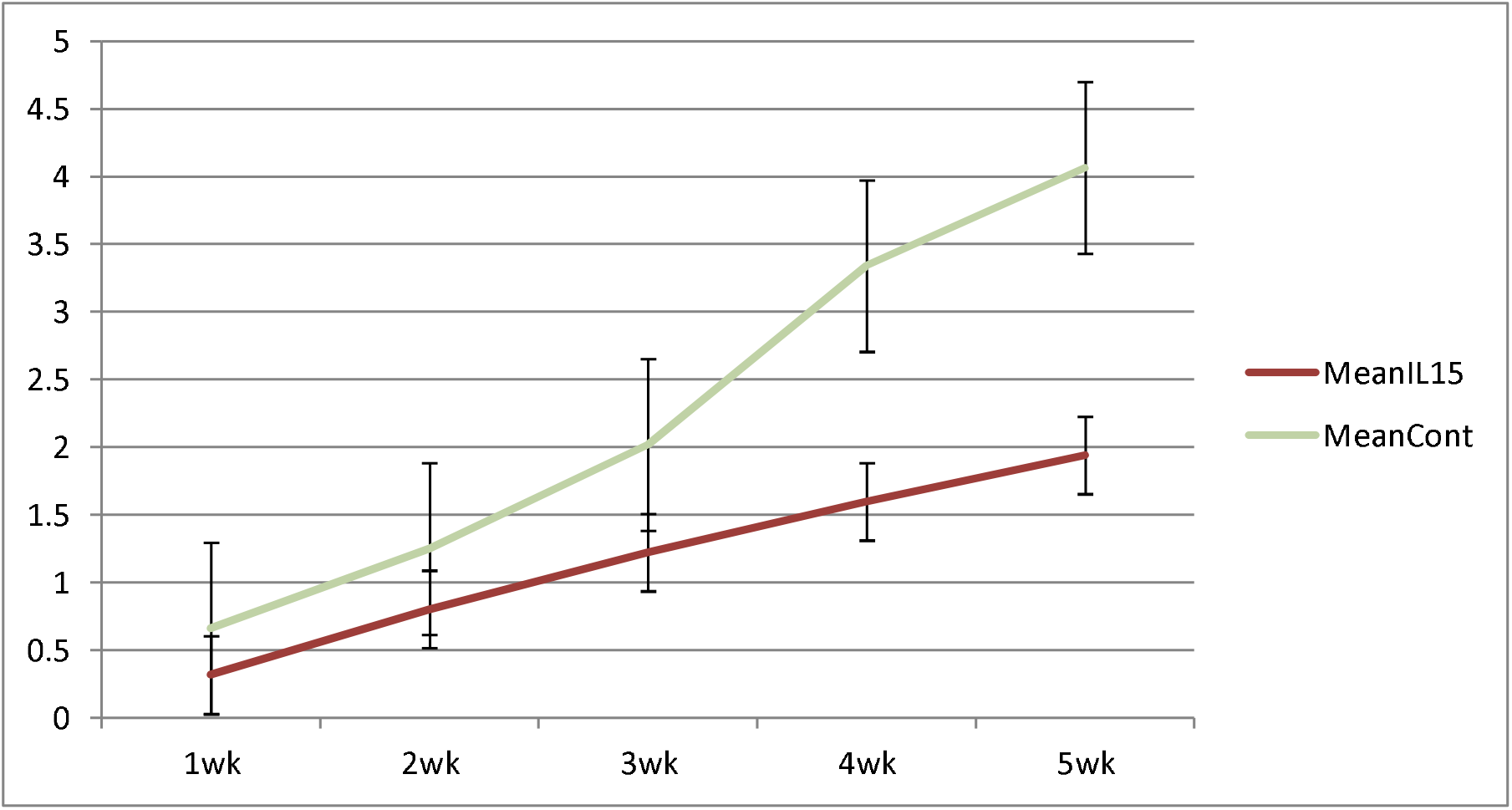
Tumor volume was measured every week after 4T1 cell inoculation with or without IL-15 treatment. Green line means control group and red line means the IL-15 group. IL-15 therapy caused statistically significant tumor volume shrinkage(standard error is used in the graph)

**Figure 2-2:**
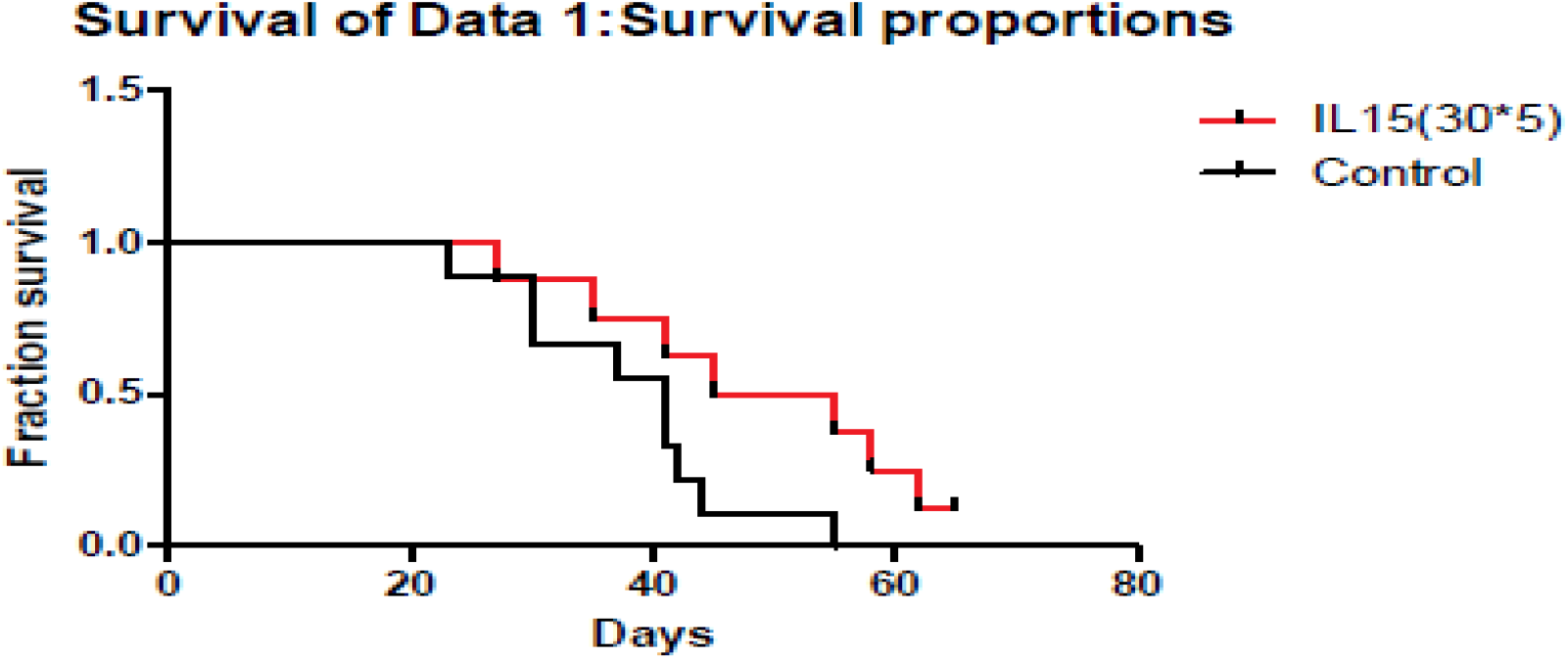
Mice survival was measured each week after 4T1 cell inoculation with or without IL-15 treatment. Red line means IL-15 treated group and black line means control group. IL-15 treatment significantly prolonged mice survival. (Likelihood P value=0.048, median survival for control group=41 days and for IL-15 treated group=50 days)

### IFN-γ enhances tumor volume and reduces mice survival in 4T1 model

After an IFN-γ IP injection (50000IU x 5days), surprisingly, we find that IFN-γ enhances tumor size compared to that of the control mice. In addition, IFN-γ injection reduces the mice survival rate compared to that of the control mice (Figure 3). IFN-γ has a negative effect on immunity against tumors.

**Figure 3-1:**
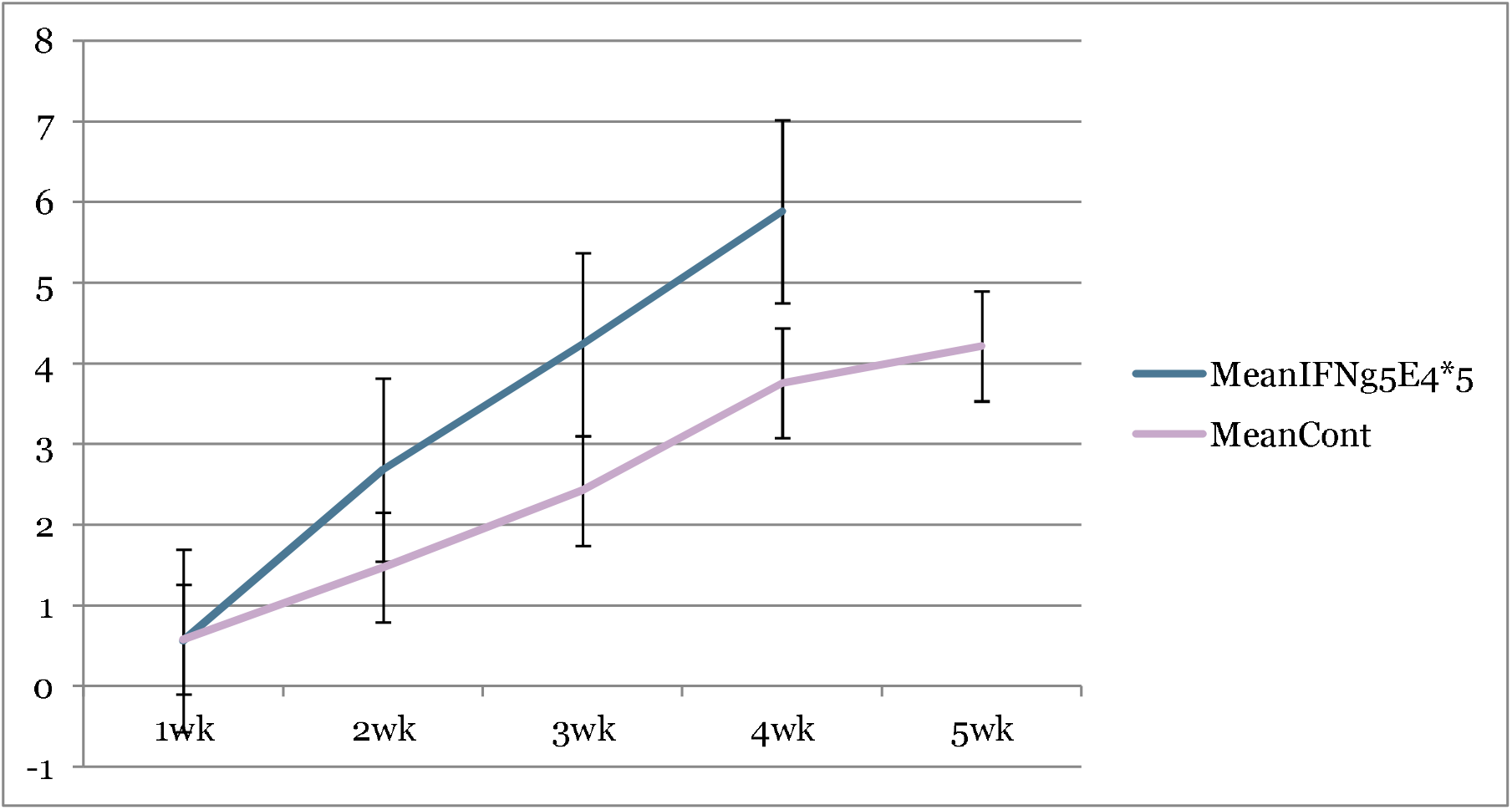
Tumor volume was measured every week after 4T1 cell inoculation with or without IFN-γ treatment. Purple line means control group and blue line means the IFN-γ group. IFN-γ therapy caused statistically significant tumor volume enlargement (standard error is used in the graph)

**Figure 3-2:**
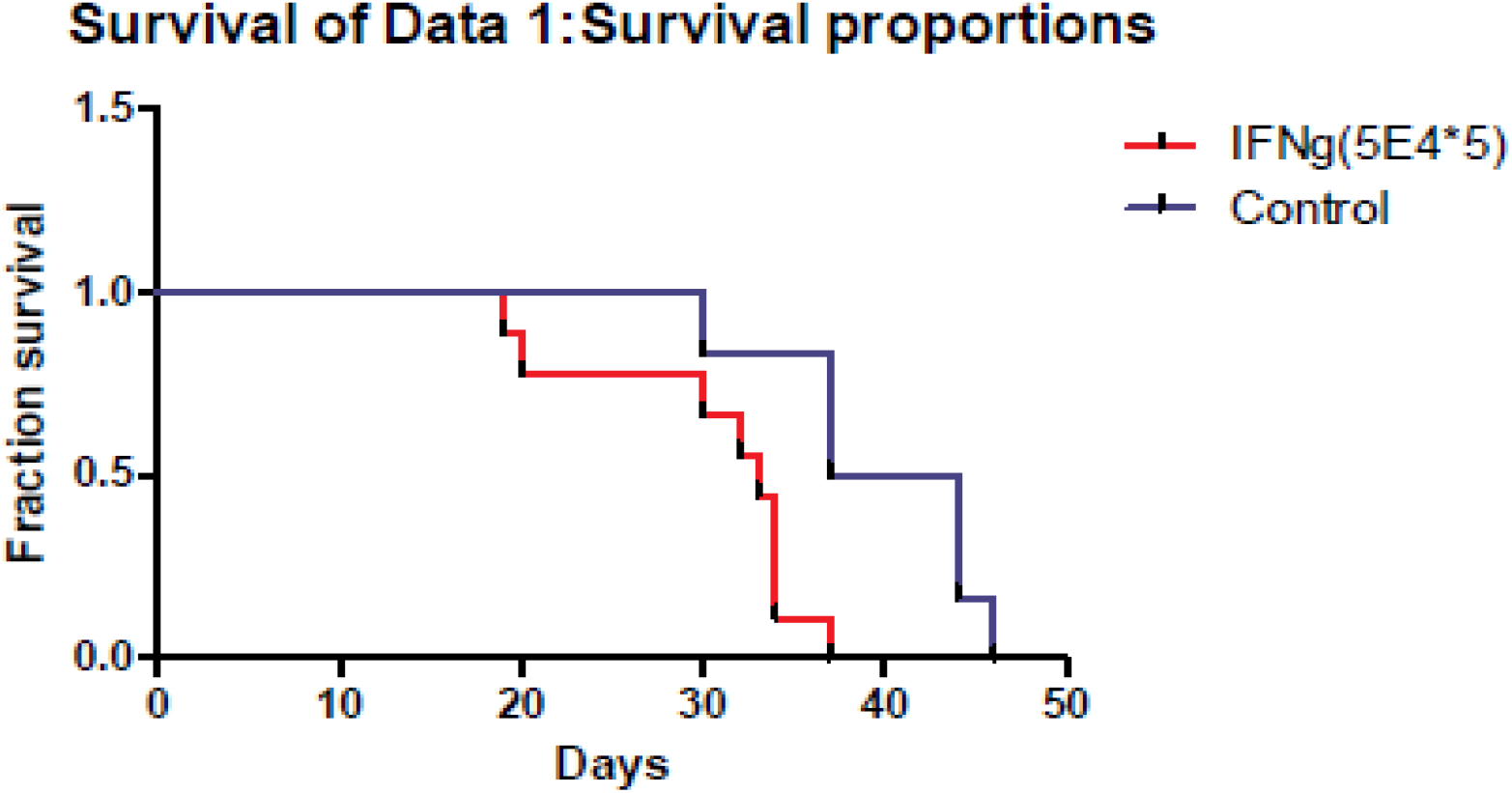
Mice survival was measured each week after 4T1 cell inoculation with or without IFN-γ treatment. Red line means IFN-γ treated group and blue line means control group. IFN-γ treatment significantly shortened mice survival. (Likelihood P value=0.01, median survival for control group=40.5 days and for IFN-γ treated group=33 days)

### IL-10 or 14G2A suppresses tumor volume and prolong mice survival in an established NXS2 cancer model

IL-10 can also suppress tumor volume and prolong mice survival in a NXS2 tumor model of AJ mice. 14G2A anti-GD2 antibody can also suppress tumor volume and prolong mice survival in NXS2 tumor model. However, a combined IL10 and 14G2A effect on tumor volume and mice survival is similar to 14G2A when used alone. The effectiveness of IL-10 alone is better than either 14G2A alone or 14G2A when combined with IL-10 treatment. It is worth noting that IL-10 alone treatment caused result in a tumor regression in three AJ mice out of 10 mice, and the three mice lived healthily even after the 4-month total follow-up period. This suggested that IL-10 treatment has a very potent anti-tumor ability compared to current cancer therapeutic agents (Figure 4).

**Figure 4-1:**
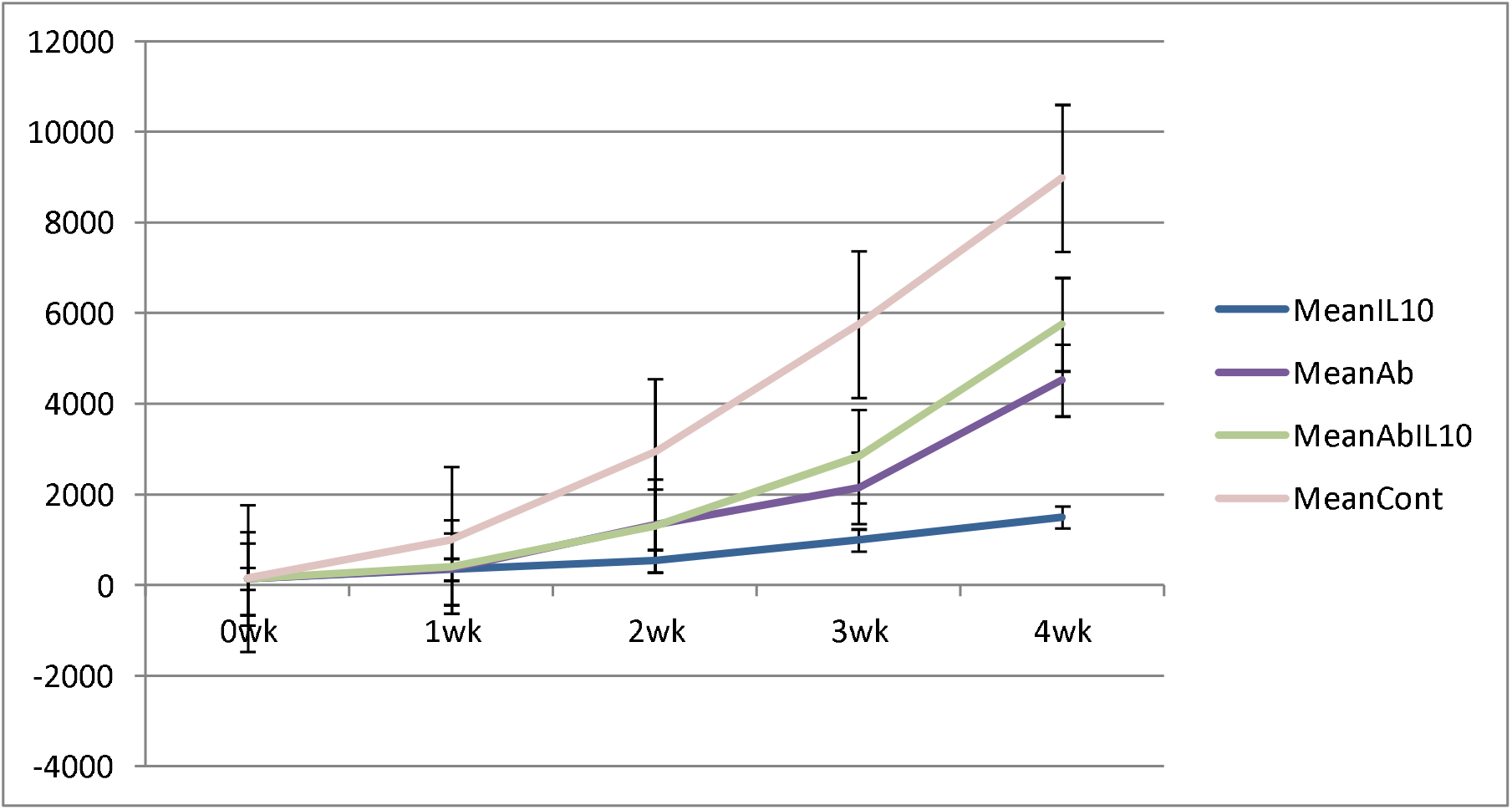
Tumor volume was measured every week after NSX2 cell inoculation with or without IL-10 treatment. Purple line means control group, blue line means the IL-10 group, dark blue line is the 14G2A antibody group, and green line is the 14G2A antibody plus IL-10 group. IL-10 therapy caused statistically significant tumor volume shrinkage. Antibody alone or antibody plus IL-10 caused mild-moderate tumor volume shrinkage. (standard error is used in the graph)

**Figure 4-2:**
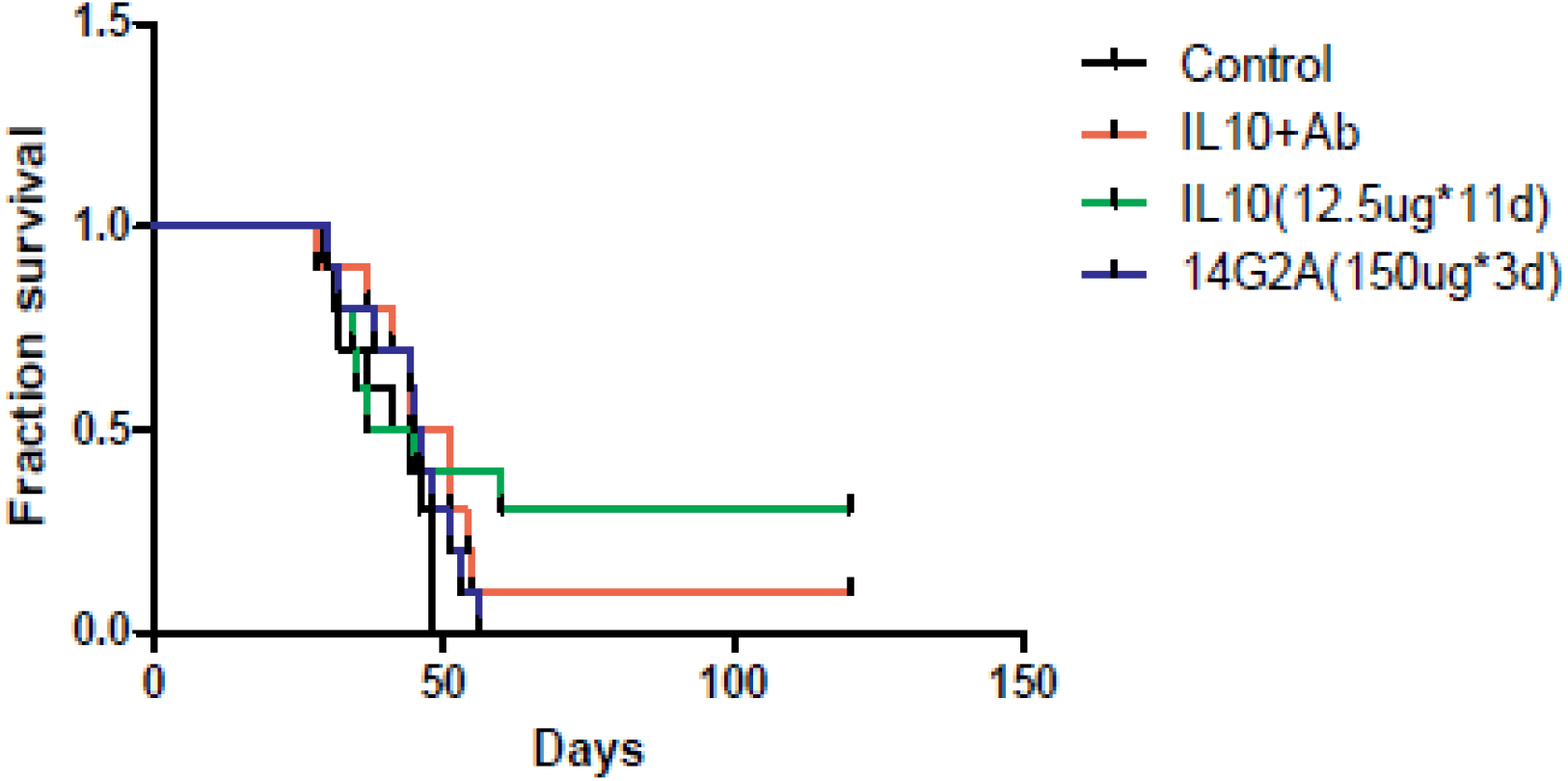
Mice survival was measured each week after NXS2 cell inoculation with or without IL-10 treatment. Green line means IL-10 treated group, black line means control group, blue line means 14G2A antibody group, and red line means 14G2A antibody plus IL-10 group. IL-10 treatment most significantly prolonged mice survival.

### IL-10 reduces cell numbers of 4T1 cells and NXS2 cells in vitro

In the pilot studies, we tested IL-10 in vitro toxicity on NXS2 and 4T1 tumor cells. After IL-10 treatment, 4T1 and NXS2 cell counts were significantly lower when compared to those without IL-10 treatment (Table 1&2). Then, we used Alamar blue assay to look at the dose response relationship of IL-10 in vitro toxicity on 4T1 and NXS2 cells (Figure7). We found that decreased 4T1 and NXS2 cell numbers depended on the use of an IL-10 concentration. A higher IL-10 dosage caused more tumor cell suppression. This effect could be seen in mouse macrophage RAW cell lines with IL-10 treatment, but IL-10 had no effect on NIH 3T3 fibroblast cell line (Figure5). It seems that IL-10 has been selective with respect to in vitro toxicity on tumor cell lines.

**Table I.** NXS2 cell counts after IL-10 treatment for three days(three replicates: A,B,C, P value=0.02, sample means are different)

| NXS2 | A | B | C | Mean |
| --- | --- | --- | --- | --- |
| Control | 680000 | 680000 | 640000 | 670000 |
| IL10<br>2ug/cc | 580000 | 660000 | 580000 | 610000 |
| IL10<br>4ug/cc | 490000 | 580000 | 560000 | 540000 |

**Table II.**
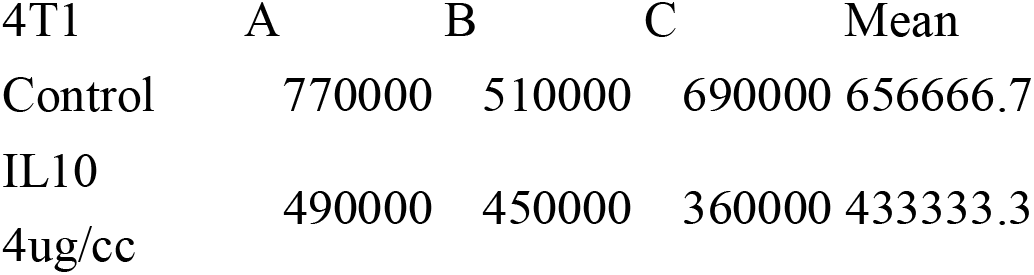
4T1 cell counts after IL-10 treatment for three days(three replicates:A,B,C, P value=0.01,sample means are different)

| 4T1 | A | B | C | Mean |
| --- | --- | --- | --- | --- |
| Control | 770000 | 510000 | 690000 | 656666.7 |
| IL10<br>4ug/cc | 490000 | 450000 | 360000 | 433333.3 |

**Figure 5-1.**
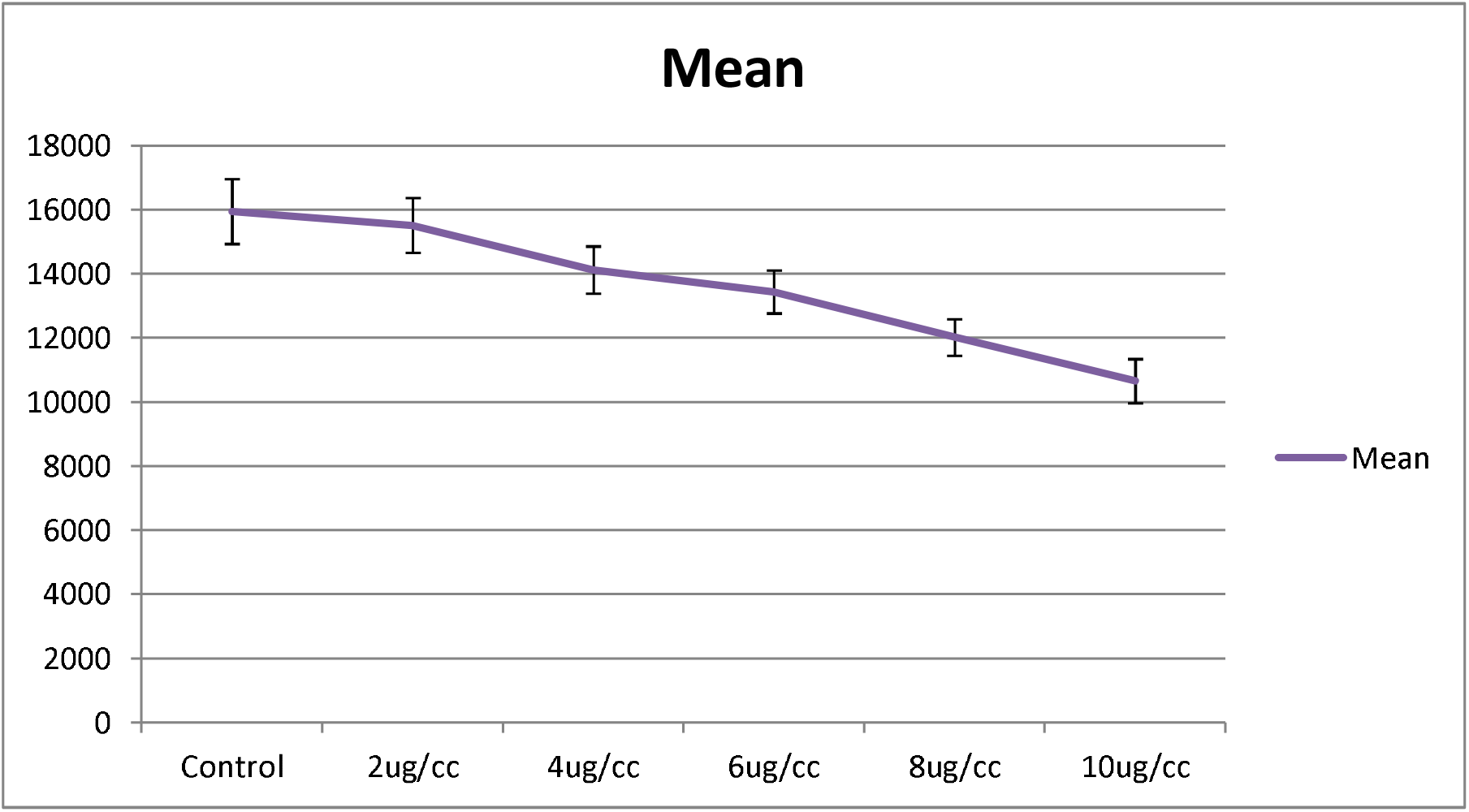
Alamar blue cell titer results of 4T1 cells treated with varies dosages of IL-10. A dose-response graph is noted and IL-10 has direct in vitro toxicity on 4T1 cells. (standard error is used in the graph)

**Figure 5-2.**
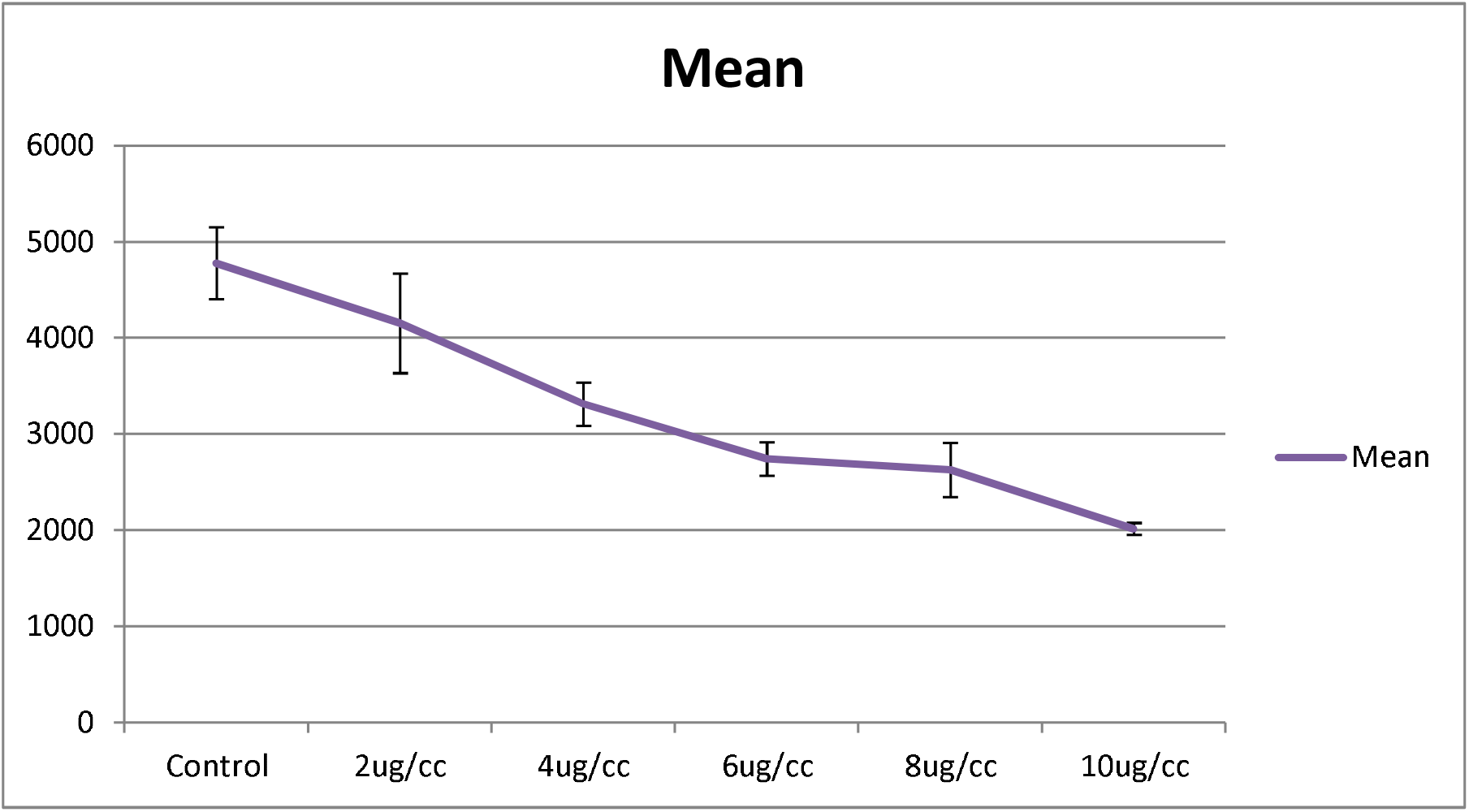
Alamar blue cell titer results of NXS2 cells treated with varies dosages of IL-10. A dose-response graph is noted and IL-10 has direct in vitro toxicity on NXS2 cells. (standard error is used in the graph)

**Figure 5-3.**
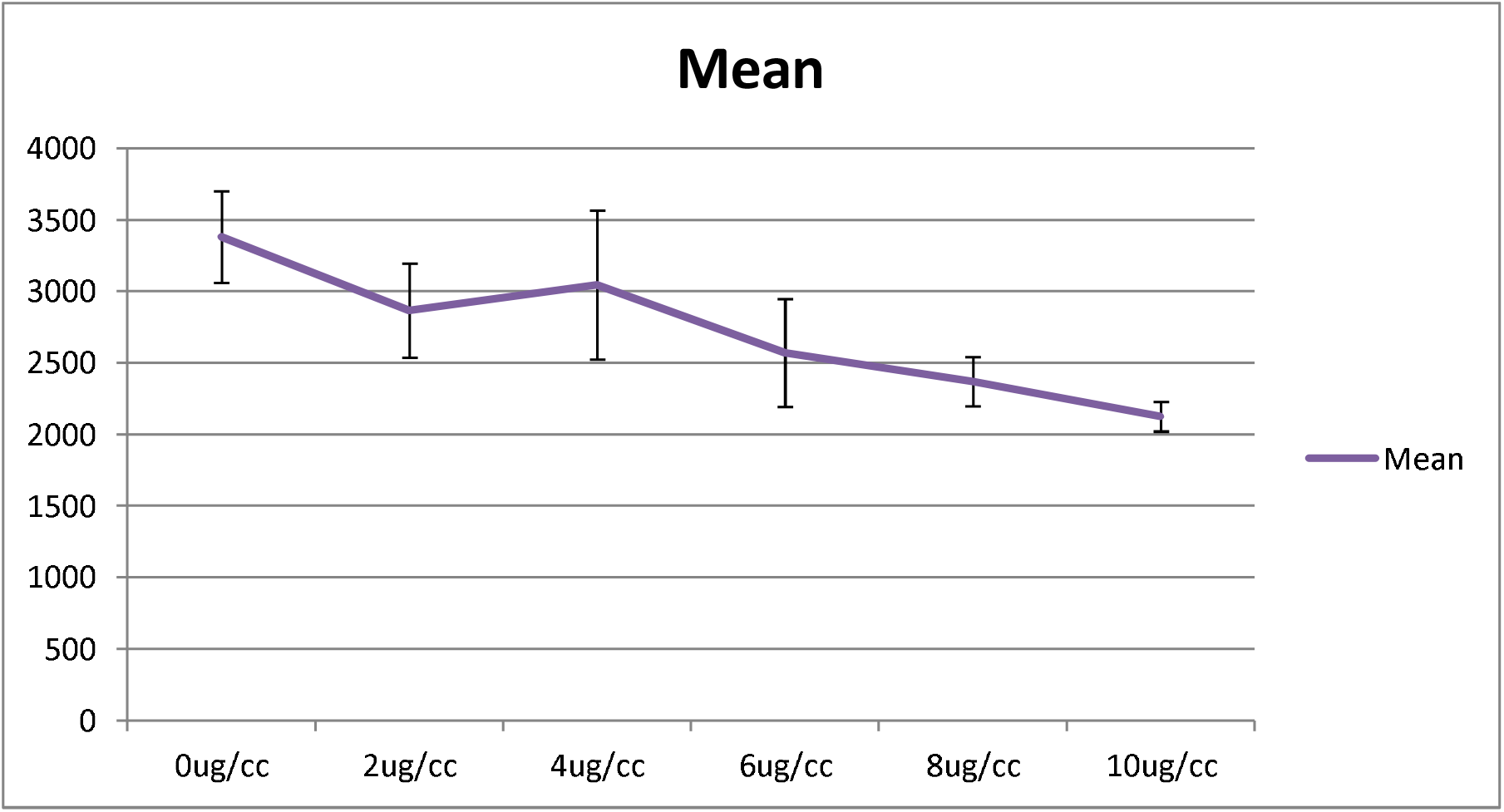
Alamar blue cell titer results of RAW macrophage cells treated with varies dosages of IL-10. A dose-response graph is noted and IL-10 has direct in vitro toxicity on RAW macrophage cells.(Positive control) (standard error is used in the graph)

**Figure 5-4.**
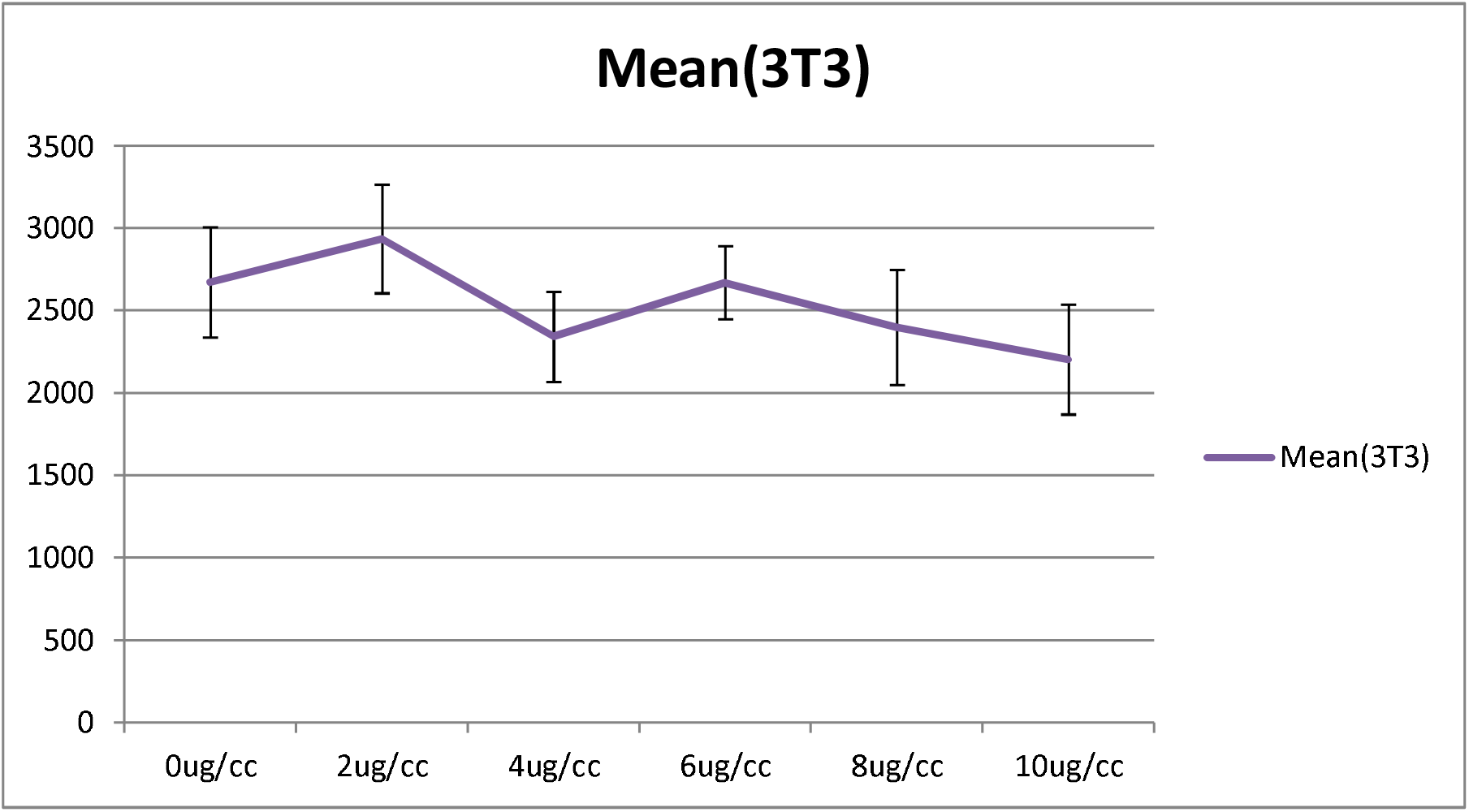
Alamar blue cell titer results of NIH 3T3 fibroblast cells treated with varies dosages of IL-10. No dose-response graph is noted and IL-10 has no in vitro toxicity on NIH 3T3 fibroblast cells.(Negative control) (standard error is used in the graph)

### Fc receptor (CD16/32/64) and IL-10 receptor (CD210) expression on NXS2 and 4T1 cells

We used flowcytometry to check Fc receptor (CD64, CD16 & CD32) and IL-10 receptor expression on NXS2 cells and 4T1 cells. There is neither CD16/ CD32 nor CD64 expression on either NXS2 cells or 4T1 cells. Besides, IL-10 treatment could not stimulate the expression of Fc receptors on NXS2 tumor cells. Surprisingly, IL-10 receptor expression was found both in 4T1 cells and NXS2 cells. And, there were two subpopulations of 4T1 or NXS2 cells with different levels of IL-10 receptor expression. Thus, IL-10 might have a direct inhibition effect on 4T1 and NXS2 cells via their IL-10 receptor (Figure 6 and 7).

**Figure 6-1.**
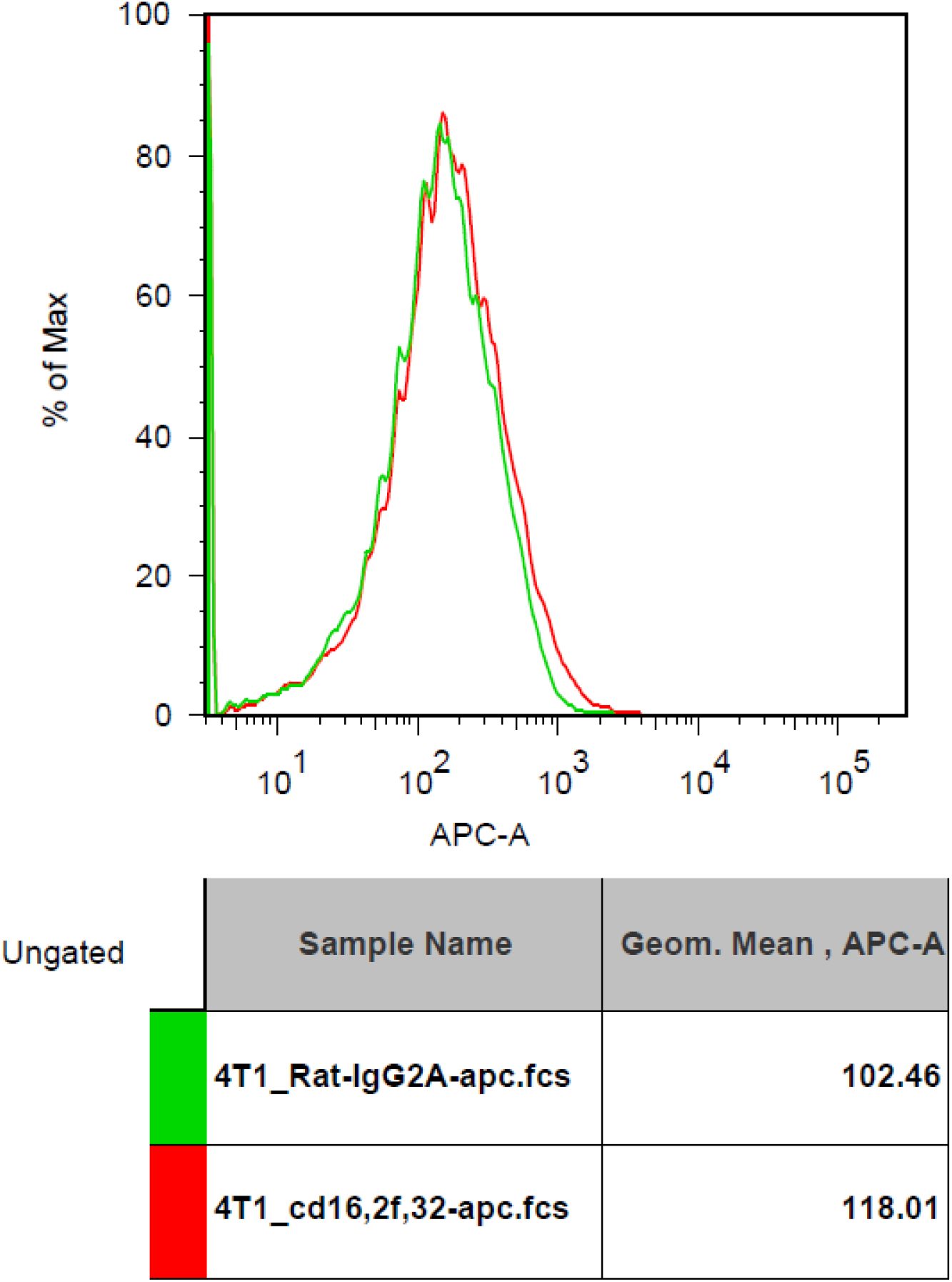
CD16/32 Fc receptor expression on 4T1 cells compared to isotype control. There was no CD16/32 expression on 4T1 cells.

**Figure 6-2.**
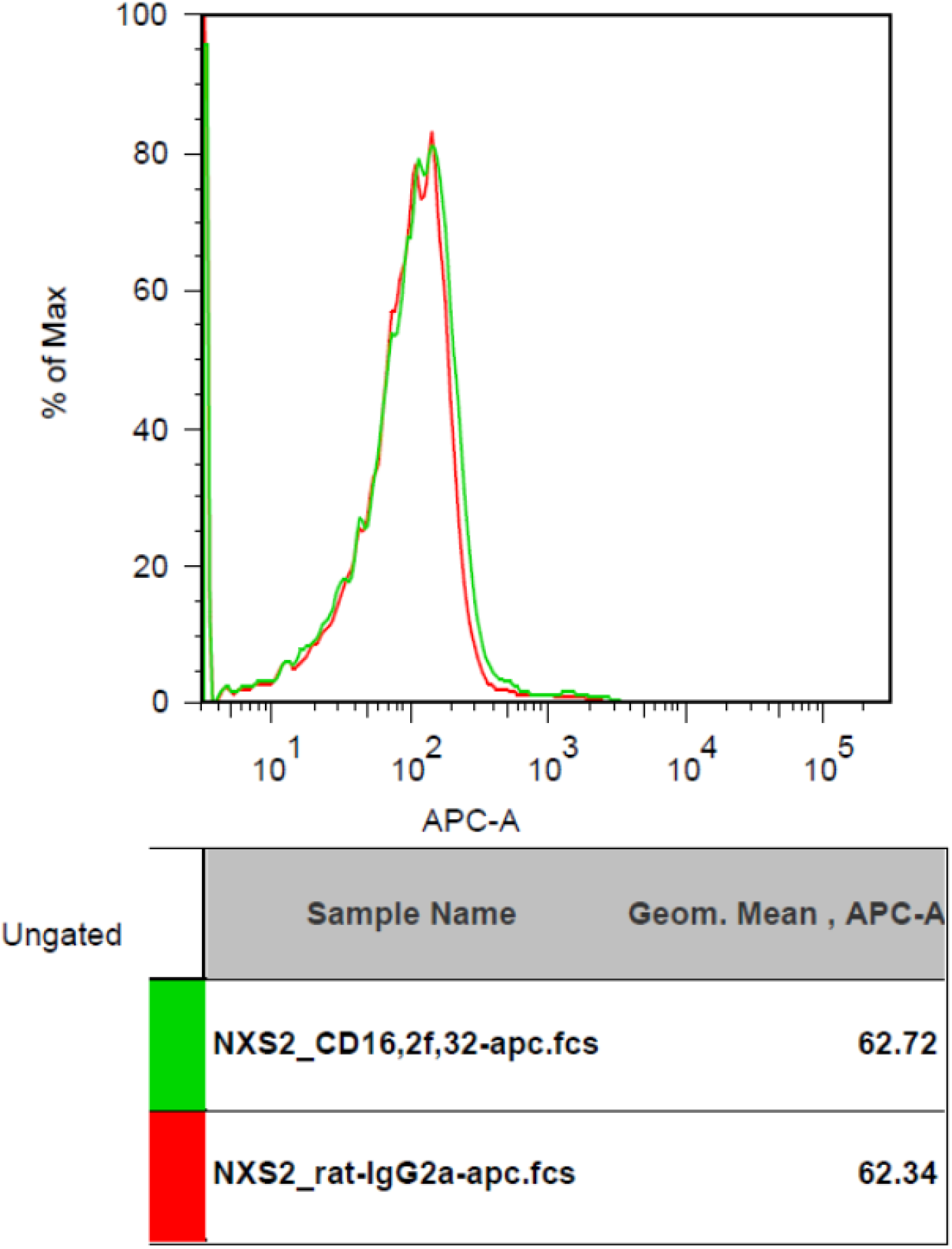
CD16/32 Fc receptor expression on NXS2 cells compared to isotype control. There was no CD16/32 expression on NXS2 cells.

**Figure 6-3.**
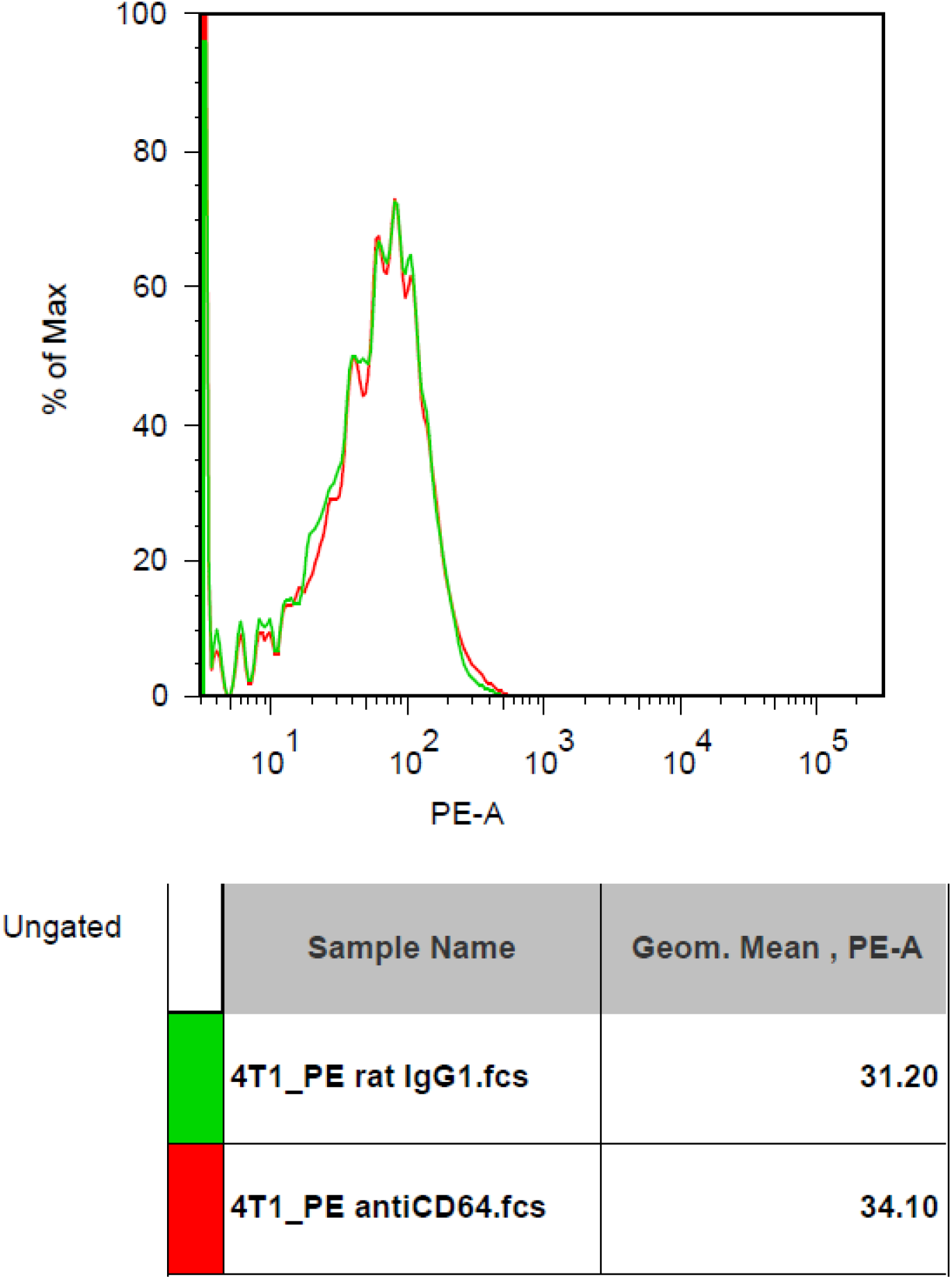
CD64 Fc receptor expression on 4T1 cells compared to isotype control. There was no CD64 expression on 4T1 cells.

**Figure 6-4.**
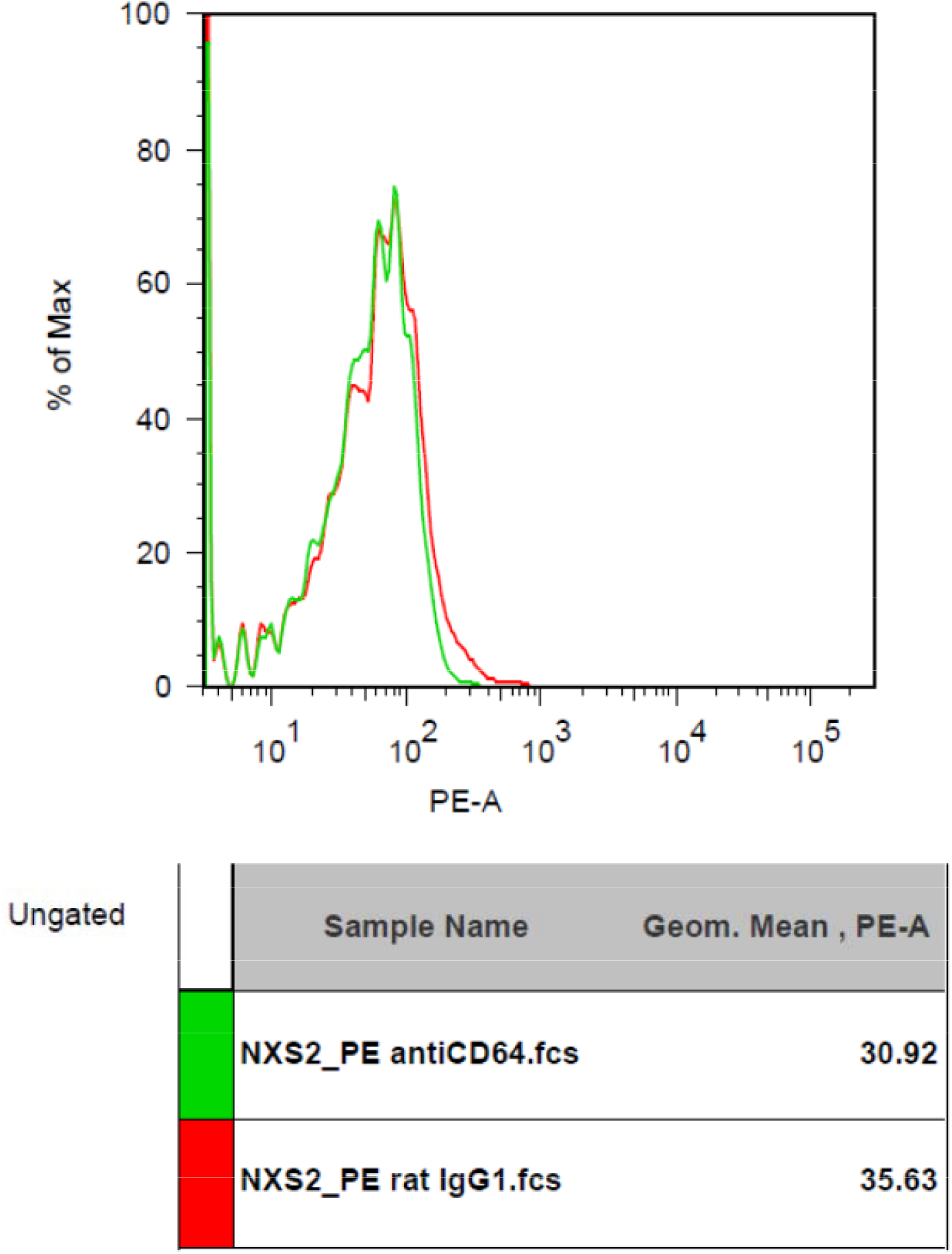
CD64 Fc receptor expression on NXS2 cells compared to isotype control. There was no CD64 expression on NXS2 cells

**Figure 6-5.**
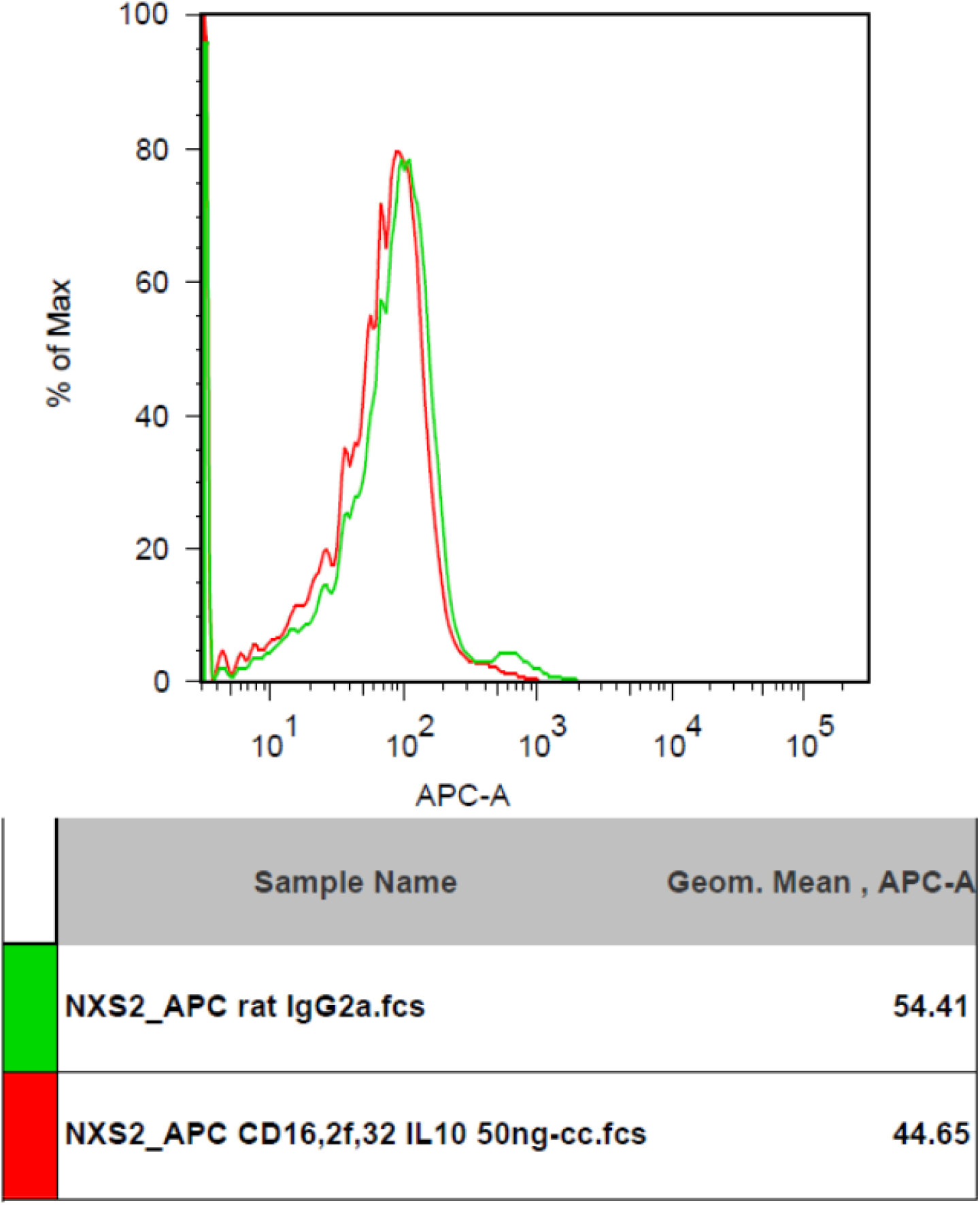
CD16/32 Fc receptor expression on NXS2 cell treated by IL-10 50ng/cc for two days. There was no expression of CD16/32.

**Figure 6-6.**
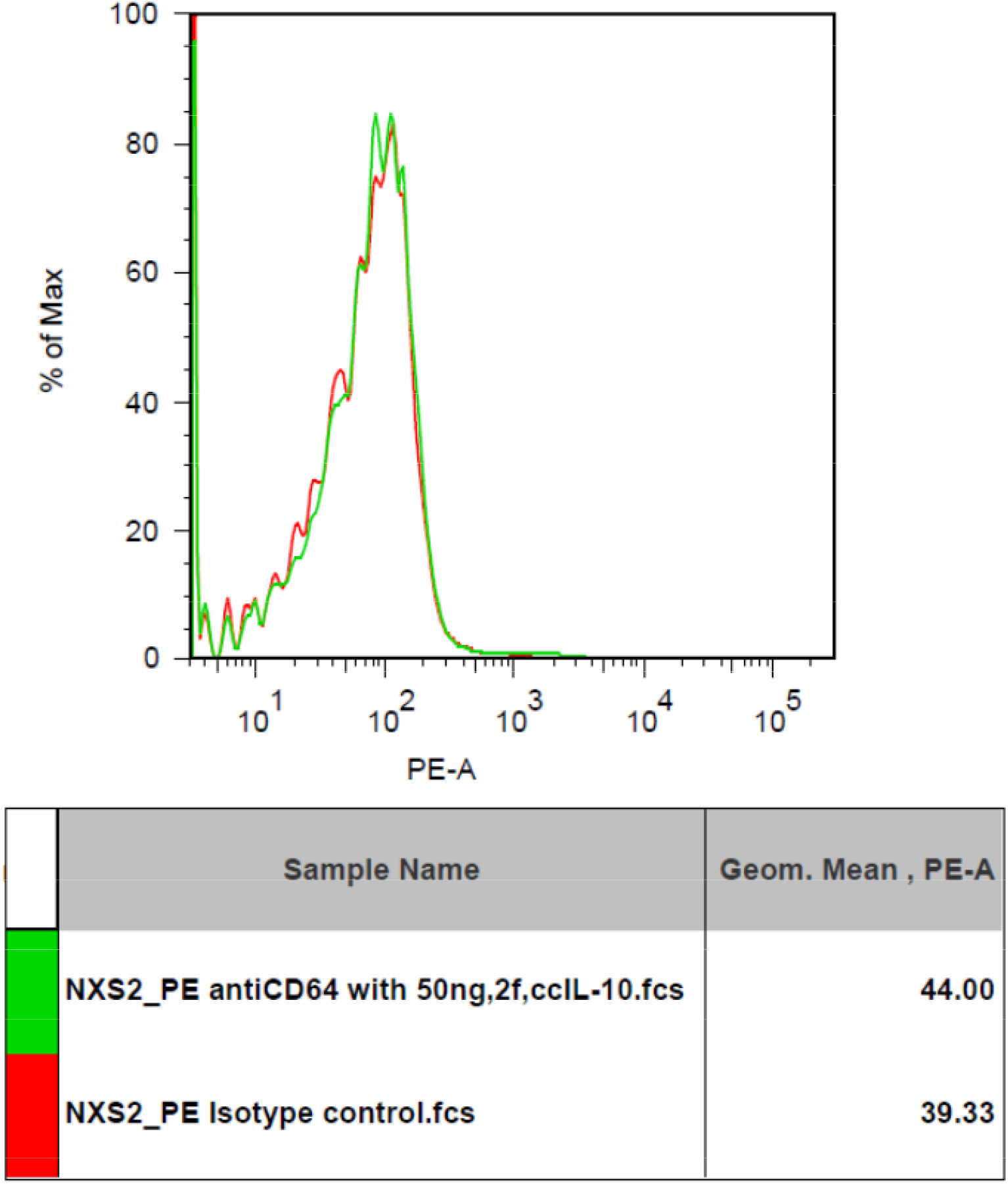
CD64 Fc receptor expression on NXS2 cell treated by IL-10 50ng/cc for two days. There was no expression of CD64.

**Figure 7-1.**
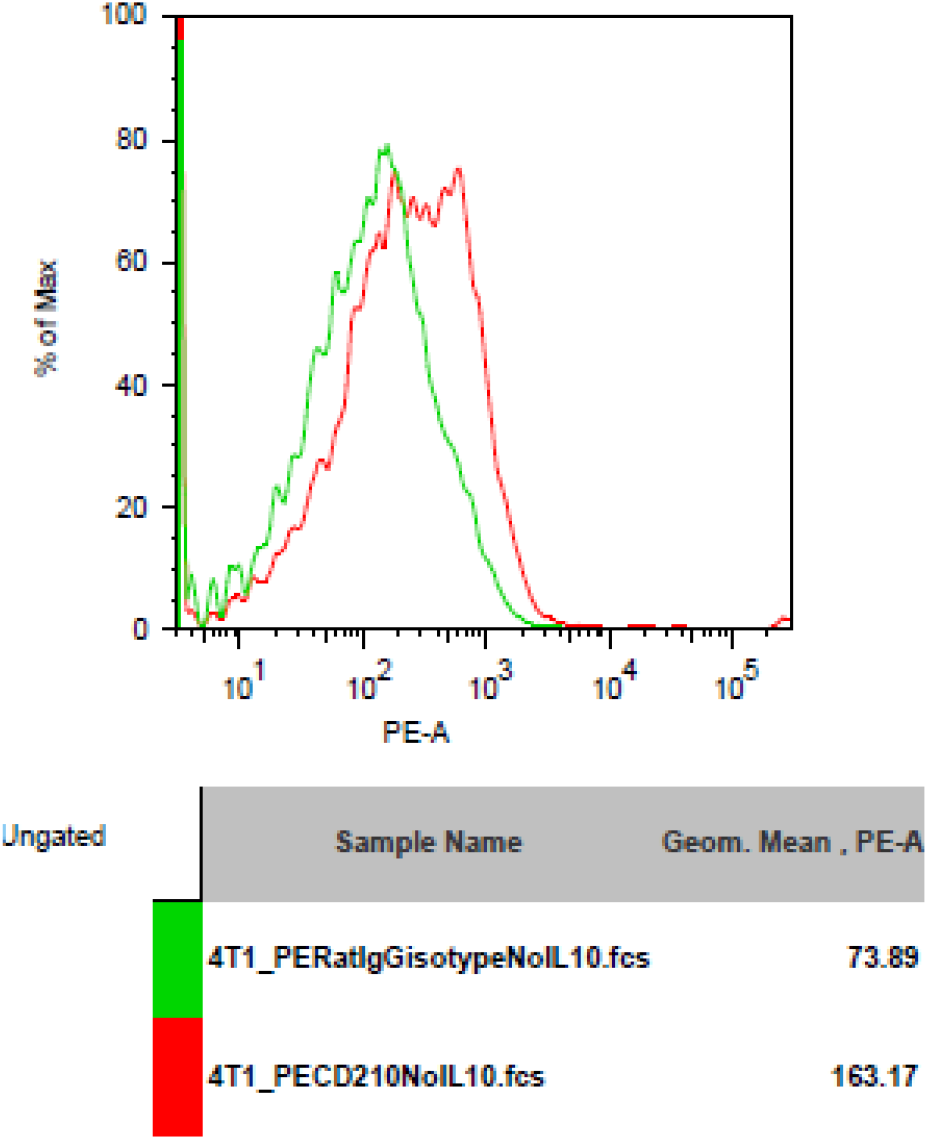
IL-10 receptor(CD210) expression on 4T1 cells. There was strong CD210 expression on 4T1 cells compared to isotype control.

**Figure 7-2.**
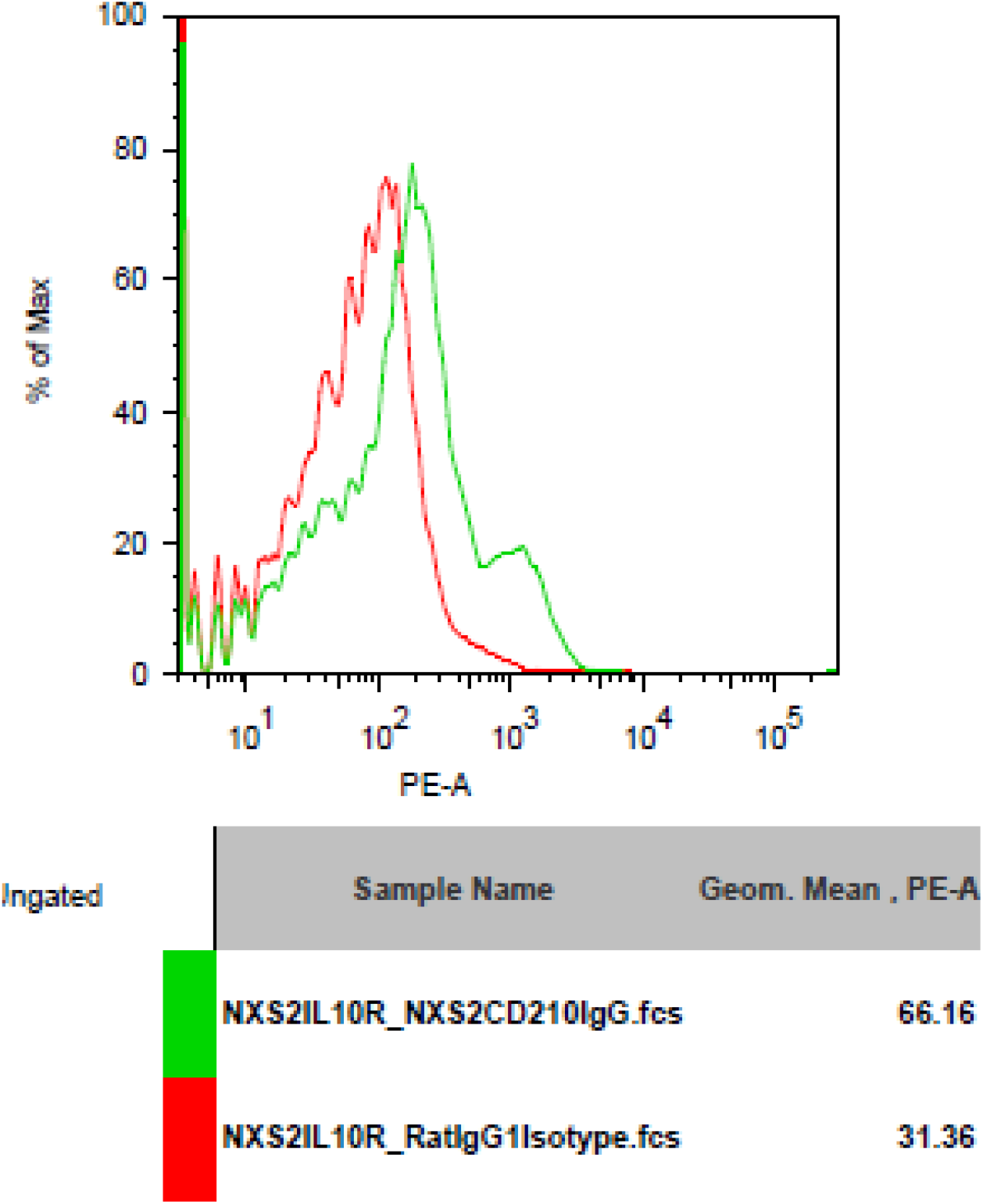
IL-10 receptor(CD210) expression on NXS2 cells. There was strong CD210 expression on NXS2 cells compared to isotype control.

### GD2 expression induction in immune cells after IL-10 treatment

GD2 antigen expression level was assessed after IL-10 treatment in spleenocytes of Balb/c mice. We found that IL-10 can induce GD2 antigen expression on immune cells. We found that GD2 antigen is expressed on spleenocytes in control mice. In addition, IL-10 treatment for one or three days enhances the expression of GD2 on spleenocytes (Figure 8).

**Figure 8-1.**
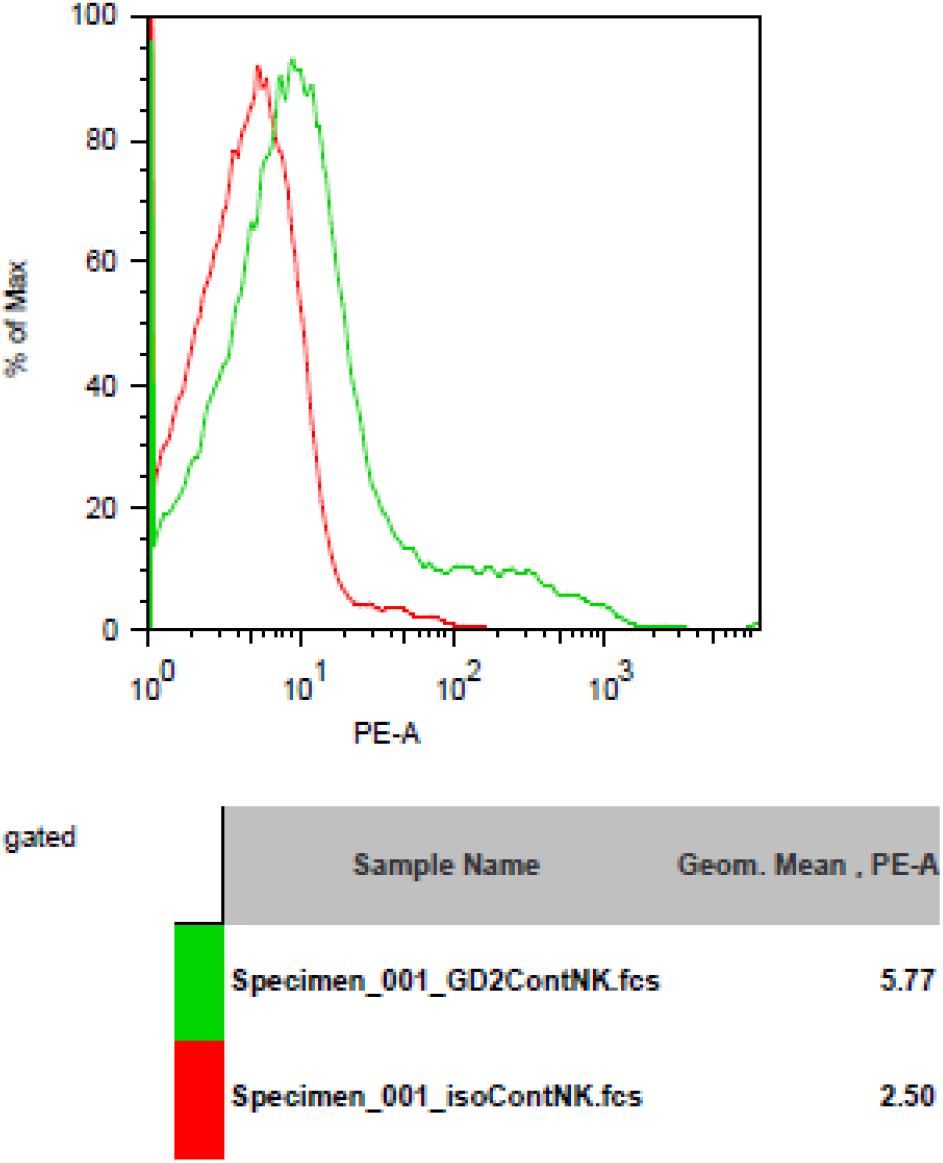
GD2 antigen expression on spleenocytes from normal mice compared to isotype control. There was GD2 expression on spleenocytes of normal mice.

**Figure 8-2.**
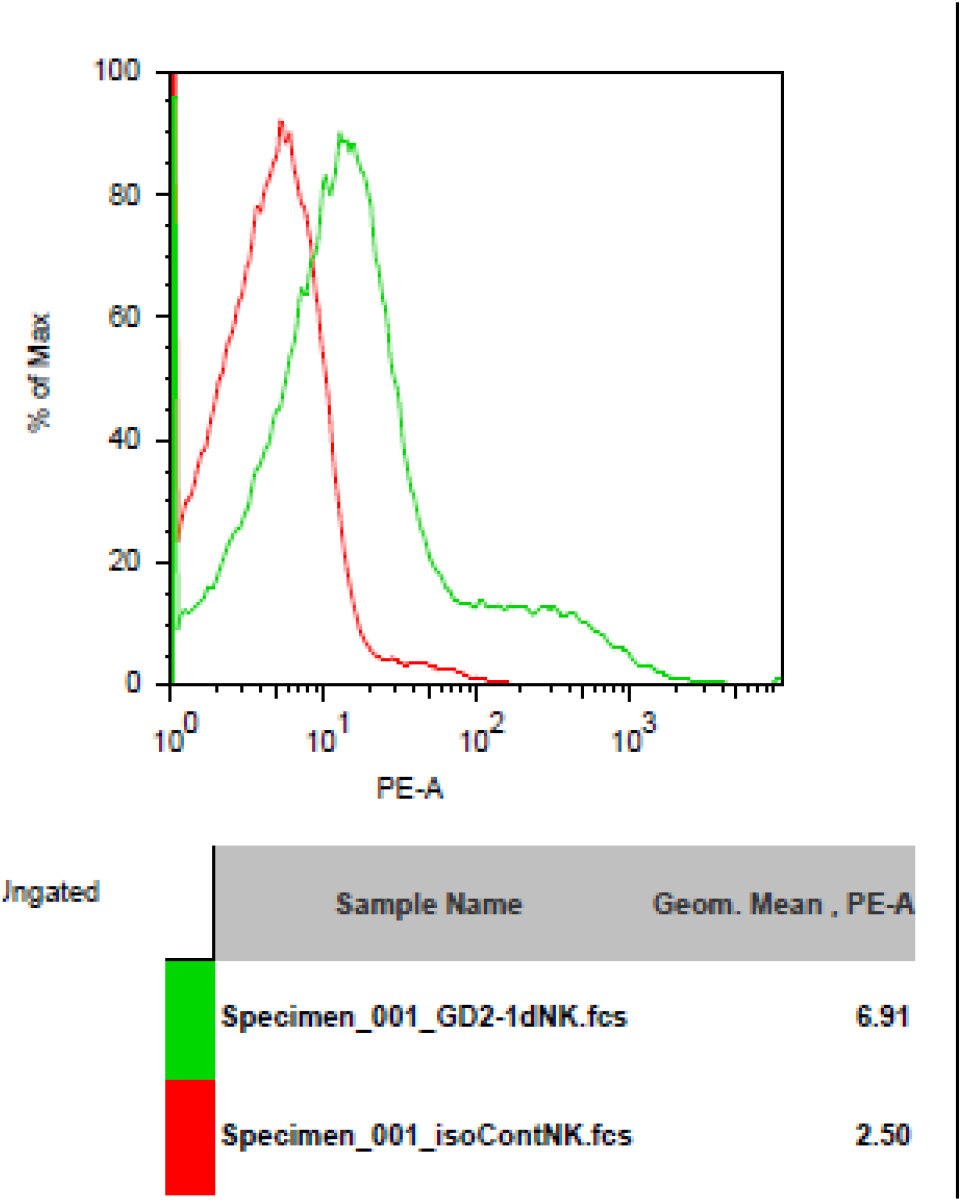
GD2 antigen expression on spleenocytes from mice injected with IL-10 for one day compared to isotype control. There was GD2 expression on spleenocytes of mice with one dose of IL-10. The magnitude of GD2 expression of one-dose IL-10 injected mice is greater than that of normal mice.

**Figure 8-3.**
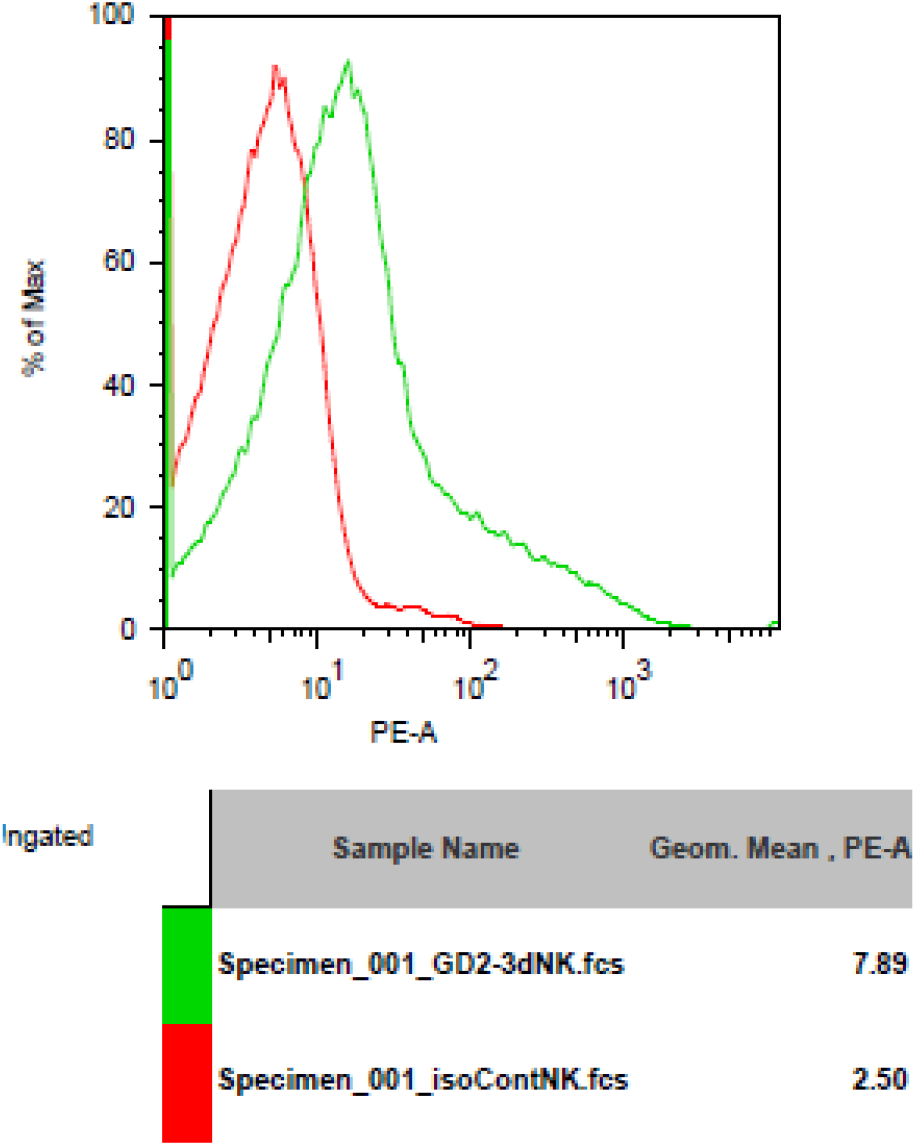
GD2 antigen expression on spleenocytes from mice injected with IL-10 for three days compared to isotype control. There was GD2 expression on spleenocytes of mice with three doses of IL-10. The magnitude of GD2 expression of three-dose IL-10 injected mice is greater than that of normal mice and than that of one-dose IL-10 injected mice.

## Discussion

THαβ immunity is a newly proposed host immunological pathway. Thus, four types of host immune responses can match four types of hypersensitivities (1–3). Traditional TH1 is a macrophage dominant delayed-type hypersensitivity. TH2 is both an eosinophil and basophil dominant IgE mediated hypersensitivity. TH17 is a PMN centered immunoglobin complex mediated hypersensitivity. And, the THαβ immunity is an antibody dependent cellular cytotoxicity hypersensitivity. In THαβ immune response, NK cell antibody dependent cellular cytotoxicity is the major mechanism which can kill virus infected cells. Thus, we can also use this mechanism to kill tumor cells with tumor antigens present in the cell surface. Other immunological pathways are not suitable for cancer immunotherapy. TH2 immune response use histamine-mediated physiological expulsion to expel parasites. It cannot be used for cancer immunity. TH17 with key mediator-TGFβ is not only a molecule for angiogenesis but also a molecule for suppressing immunity by activating Treg cells. TH17 can even enhance tumor growth. In traditional TH1 immunity, macrophages play important roles. And, macrophages enhance tumor invasiveness and metastasis. Traditional TH1 immunity is not a good technique for cancer immunotherapy. THαβ immunological pathway is the best candidate.

THαβ immunity includes IL-10(+++)/IFN-γ(+) CD4 T cells(3). IL-10 plays a central role in mediating a THαβ immunological pathway. Thus, we deduce that IL-10 plays a key role in initiating an NK cell ADCC response. This conclusion was suggested in a previous in vitro study. Our study did find out that IL-10 has significant effect on tumor shrinkage and mice survival. In previous studies, an IL-10 expressing tumor can mediate an immunity rejecting tumor. IL-10 can also reduce the metastasis of cancer in a mice model(8, 9). According to these results, we highly suggest the use of IL-10 for cancer immunotherapy. Previous studies have controversial evidences about IL-10’s anti-tumor ability. Some papers suggest that IL-10 can suppress a tumor. However, other papers suggest that tumor associated IL-10 elevation can suppress a hosts anti-tumor immunity(10, 11). Our findings support the first hypothesis. IL-10 has a strong anti-tumor ability. In clinical cancer patients, IL-10 may be up-regulated to initiate anti-cancer immune response. However, the elevation level of IL-10 may not high enough to combat cancer. Thus, these authors had incorrect explanations about an IL-10’s role in tumor immunity.

In a previous in vitro study, IL-15 worthy can activate NK cell/ T cell and has anti-tumor immunity (12, 13). In viral infection, an IL-15 is usually up-regulated to induce NK cells and memory T cells proliferation. In this research, we also find that an IL-15 has significant effect on tumor shrinkage and on prolong mice survival. Although the effectiveness of IL-15 is not an impressive as IL-10, we also suggest the use of IL-15 for cancer immunotherapy. IL-15 belongs to the IL-2 cytokine family. In a previous study, IL-15 has a wider therapeutic window with fewer side effect compared to IL-2. IL-15 is also a good candidate for cancer treatment.

IFN-γ is usually considered to be a medication for cancer therapy. However, up to now, IFN-γ treatment in cancer clinical trials only had limited success. Some reports even found that IFN-γ can worsen the prognosis of cancer patients. Surprisingly, we found that IFN-γ can reduce mice survival with an enlarged tumor size compared to the control mice. IFN-γ is a major TH1 cytokine which can be used to activate monocytes/macrophages. In several previous animal studies, murine IFN-γ can increase cancer metastasis in mice(14, 15). Because IFN-γ is a strong macrophage activator and because IFN-γ can induce macrophage fusion, we suggest that IFN-γ could be harmful in cancer treatment. Cancer metastasis is thought to be mediated by tumor-macrophage fusion(16). After tumor-macrophage fusion, cancer cells are brought to macrophage reservoir pools in our body such as an individual brain(microglia), liver(Kupffer cells), spleen(spleen macrophages), bone(osteoclast), and lung(alveolar macrophages). Tumor-macrophage fusion can also enhance angiogenesis and cancer invasiveness because a tumor can acquire Toll-like receptors from monocytic cells and proteases such as MMPs to disrupt an extracellular matrix(17–19). Thus, it is reasonable in our results that IFN-γ is detrimental to host immunity against cancer. IL-10 receptor is expressed on hematopoietic cells including T cell, B cell, NK cell, mast cell, dendritic cell, and monocyte/macrophage. In addition, Fc receptors are usually expressed terminal differentiated macrophages(20). Thus, our results implied that cancer-macrophage fusion is actually a cancer-monocyte fusion. Fusion with terminal differentiated macrophage can stop cell proliferation(21). Besides, IL-10 is harmful to cancer cells and evolution process cannot let cancer to express IL-10 receptor to harm itself. It implies that cancer does have a monocyte origin to explain this phenomenon. The cancer cell lines from our study must originate from fused monocytic cells(22).

Our first tumor model is murine breast cancer. However, unlike human breast cancer, murine breast cancer doesn’t have Globo H antigen, so we could not use a VK9 anti-Globo H antibody to treat the 4T1 mice breast cancer model(data not shown)(5). Our second tumor model is murine neuroblastoma. Our original hypothesis is that IL-10 plus anti-GD2 antibody may activate an NK cell ADCC to kill cancer cells(6). Surprisingly, we found that IL-10 plus anti-GD2 antibody treatment is no better than IL-10 alone. IL-10 alone treatment groups have the longest survival and the smallest tumor volume. Previous references suggested that tumor cells can express Fc IgG receptor to bind and interfere antibody’s killing ability(23, 24). However, we check CD64/CD32/CD16 Fc receptor expression on 4T1 and NXS2 cells, and we didn’t see any Fc receptor expression.

A previous paper reported that GD2 expression is found on lymphocytes(25). Therefore, we checked the GD2 expression on the spleenocytes/immune cells of mice, and we found a strong GD2 expression on these immune cells. Thus, an anti-GD2 antibody could attack immune cells themselves and thus hurt anti-tumor immunity. This deduction helps to explain our results. In addition, we found that both 4T1 and NXS2 cells have a strong IL-10 receptor expression. And, IL-10 has direct in vitro toxicity on both 4T1 and NXS2 cells. Besides, IL-10 can also kill monocytes/macrophages(26). This suggests that IL-10 can directly kill tumor cells as well as tumor associated macrophages. This study provides the first evidence that IL-10 can directly kill cancer cells. Thus, IL-10 is a highly effective anti-tumor agent.

In summary, THαβ immunity is an effective method for tumor immunity. Both IL-10 and IL-15 have a significant effect on tumor volume shrinkage as well as on prolonging mice survival. However, IFN-γ appears to have negative effect on tumor immunity. Moreover, IFN-γ treatment enhances tumor growth and reduces mice survival rate. IL-10 has direct anti-tumor and anti-macrophage toxicity in vitro. This can explain why IL-10 is so effective in tumor immunity. We strongly suggest starting a clinical trial to test IL-10 anti-tumor efficacy in clinical oncology patients in the near future.

## Materials and Methods

### Cell culture

4T1 cells and NXS2 cells were obtained from ATCC. 4T1 cells were maintained in RPMI 1640 medium and NXS2 cells were maintained in DMEM medium (Gibco Inc., MA). A sub-culture was performed every three days.

### Mouse cancer model and treatment

4T1 cells and NXS2 cells were inoculated subcutaneously into the right flank of Balb/c female mice and AJ female or male mice respectively in day0. These mice were among 6-8 weeks-old and were purchased from the National Experimental Animal Center of Taiwan. In the 4T1 model, cytokines (hIL-10 20ug, mIL-15 15ug bid, IFN-γ 5×10^4^ IU from BioLegend Inc., CA) were injected intraperitoneally in mice on day 1, day 3, day 5, day 7, and day 9. In the NXS2 model, IL-10 12.5ug*10 days IP with and without 14G2A (anti-GD2 antibody, from Dr. Alice L. Yu’s lab) 150 ug IV on day 1, 5, 9 were used. Tumor volume was measured every week. In addition, mice survival rate was observed and recorded.

### Cell counting experiments

A pilot study was done to calculate the IL-10 in vitro toxicity on NXS2 cells and 4T1 cells. NXS2 cells or 4T1 cells were seeded in 5cc dishes both with and without IL-10 treatment (2ug/cc or 4ug/cc) and were incubated for three days. Then, cells were harvested and automatic cell counting was done to decide the cell number with and without IL-10 treatment.

### Alamar blue assay

96 well dishes were used for cell titer Alamar blue assay (Thermo Fisher Scientific Inc., MA). 1000 4T1 cells per 200 ul were seeded in each well for one night. Then, IL-10 with varies dosages (2ug/cc-10ug/cc) was added for two days. Then, Alamar blue was added to decide viable cell counts. Similar procedures were done for NXS2 cells (2000 NXS2 cells per well with IL-10 for 3 days), macrophages (1500 RAW cells per well with IL-10 for 2 days as positive control), and fibroblasts (1500 NIH 3T3 cells with IL-10 incubation for 2 days as negative control) to obtain Alamar blue titer.

### Flowcytometry

Flowcytometry (BD Bioscience Inc., NJ) was done to decide both Globo H antigen, Fc receptor(CD16 &CD32&CD64), and IL-10 receptor(CD210) expression on 4T1 and NXS2 cells. Antibodies were purchased from eBioscience Inc, CA. Antibody of 1 ug for the above antigens was added to 2*10^5 4T1 or NXS2 cells and incubated for 30 minutes. Then, flowcytometry was performed to check the preceding antigen expressions as compared to the isotype control.

### GD2 antigen induction

In Balb/c mice, IL-10 10ug was given an IP for 1 or 3 days. Then mice were sacrificed and spleenocytes were harvested. The GD2 antigen expression on spleenocytes/immune cells was studied by anti-GD2 IgG antibody (from Dr. Alice L. Yu’s lab) with flowcytometry compared to the control mice without any IL10 treatment. Then, the result was analyzed by FlowJo software (Tree Star, Asland, OR)

## Acknowledgements

We are very thankful for the help of Dr. Felix Hung, Mr. Po-Kai Chuang, Dr. Sherry Tsui-Ling Hsu, Dr. Tai-Na Wu, Dr. Jacky JC Wu, Dr. Alice L. Yu, and Dr. Michael Hsiao’s kind help and advices. In addition, we really appreciate the high peak research project funding of the Taiwan government.

## Figure legends

Figure 1. 4T1 tumor volume and mice survival graph after IL-10 injection compared to control mice.(n=12 each group)

Figure 2. 4T1 tumor volume and mice survival graph after IL-15 injection compared to control mice.(n=9 each group)

Figure 3. 4T1 tumor volume and mice survival graph after IFN-γ injection compared to control mice (n=9 each group).

Figure 4. NXS2 tumor volume and mice survival graph after IL-10 injection compared to control mice(n=10).

Figure 5. IL-10 in vitro toxicity on cell lines shown by alamar blue cell titer assay.

Figure 6. CD16/32 expression on 4T1 or NXS2 cells

Figure 7. IL-10 receptor expression on 4T1 or NXS2 cells

Figure 8. GD2 antigen expression on spleenocytes of mice

